# Mechanical feedback drives asynchronous cell divisions during embryogenesis

**DOI:** 10.1101/2023.11.29.569235

**Authors:** Abdul N. Malmi-Kakkada, Sumit Sinha, D. Thirumalai

## Abstract

The initial stage of Zebrafish morphogenesis is characterized by a synchronous to asynchronous transition (SAT) in cell divisions. The cells divide in unison in the synchronous phase, unlike in the asynchronous phase. Despite the widespread observation of SAT in multiple organisms, there is no theoretical framework to predict the critical number of cell cycles *n*^*^ that marks the beginning of asynchronous division. Here, by probabilistically modeling cell cycle progression under the assumption that the distribution of cell division times is broadened, we predict *n*^*^ and the time at which the SAT occurs. The theory, supplemented by agent-based simulations, supports the hypothesis that the SAT emerges as a consequence of biomechanical feedback on cell division. Our results are in excellent agreement with the experiments while also explaining the cell cycle lengthening that arises as a result of biomechanical feedback. The emergence of the asynchronous phase is due to increasing fluctuations in the cell cycle times with each round of cell division. We also make several testable predictions that further sheds light on the role of biomechanical feedback in growing multicellular systems, such as during tissue and tumor growth.

---

Time-dependent synchronous behavior is pervasive across biological systems. Examples include networks of neurons^1,2^, insulin-secreting pancreatic cells^3^, to the chirping of crickets^4^, and the synchronous flashing of fireflies^5,6^. Here, we focus on a particularly fascinating example of tissue-scale synchronous behavior observed during Zebrafish development when rapid and synchronous cleavage divisions subdivide the fertilized egg into a large population of nucleated cells^7,8^. Such fast and synchronized cell divisions are observed during the early development of multiple organisms. For instance, a frog egg divides into 37,000 cells in ∼43 hours, while the Drosophila embryo proliferates to 50,000 cells in ∼12 hours, both characterized by early stages of synchronous cell divisions^8,9^.

The initial observation of spontaneous synchronization dates back to the 17*th* century when Christiaan Hyugens reported the synchronization of pendulums due to the transfer of energy. However, the reverse process of how asynchronous behavior emerges in a synchronous system is highly relevant to understanding the morphogenesis of living systems as we show here. Because cell collectives generate geometric order driven by cell divisions^10^, understanding the transition between synchronous to asynchronous cell division could provide important insights. By analyzing experimental data in light of theory and simulations, we show that the synchronous to asynchronous transition (SAT) during zebrafish morphogenesis provides vital clues on how cells integrate biochemical and mechanical signals to regulate cell division.

Regulation of cell division especially during rapid cell proliferation at early stages of development^11,12^ is necessary for robust tissue growth and morphogenesis. Biochemically, this occurs through chemical factors that undergo cyclical changes during the cell cycle^13^. Together with biochemical switches, variations in the forces experienced by cells and the mechanical properties of the microenvironment are now understood to be important in cell cycle regulation^14–16^. In particular, rapid cell divisions could lead to the build up of mechanical pressure which in turn functions as a feedback signal that limits and delays further cell divisions^17^. We show that the SAT transition serves as a powerful example to dissect the contribution of mechanical forces in controlling cell division. We uncover that the pressure experienced by cells due to mechanical interaction with its neighbors *quantitatively* explains the onset of SAT. Our prediction of cell cycle lengthening, critical for subsequent development, as a consequence of mechanical feedback on cell division further supports our hypothesis that mechanical feedback is the key driver of SAT.

## Experimental Motivation

Our study is motivated by experimental data on the time-dependent cell division patterns during the early stages of Zebrafish morphogenesis. We start with a brief description of the experiment^18^, which tracks the number of cell nuclei (a proxy for the number of cells), *N* (*t*), as a function of time (*t*) using light-sheet microscopy (Fig. 1a). Cell divisions occur at the animal pole of the Zebrafish embryo (see inset showing three distinct division cycles), roughly every ∼ 20 mins^7,18^.

**FIG. 1:**
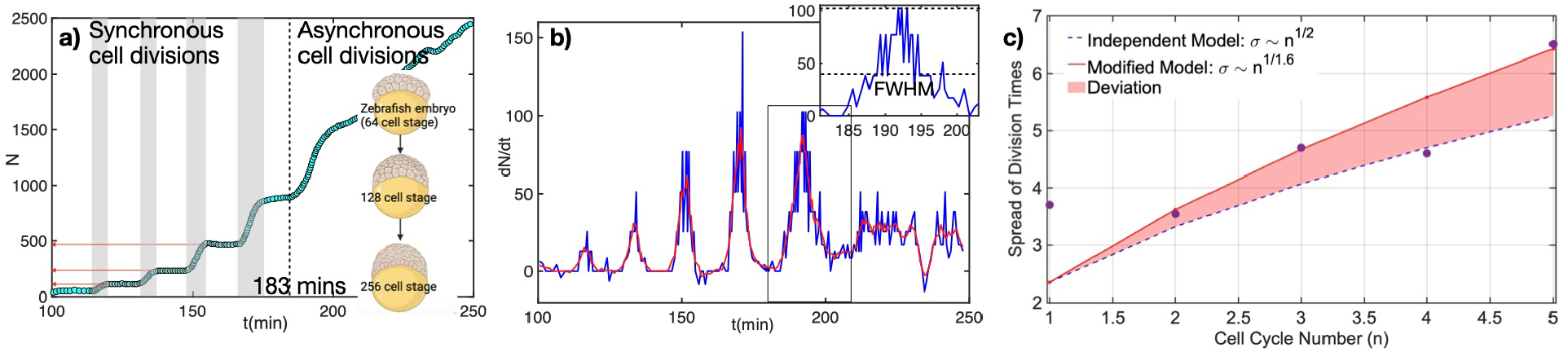
Synchronous to asynchronous transition in cell division during embryo development. **(a)** Experimental data for the number of nuclei in the Zebrafish embryo animal hemisphere from 64-cell stage to 2,000+ cells. Initially, cell divisions are highly synchronized (shaded areas) exhibiting a staircase-like pattern of number growth. After 5 cell division rounds, the asynchronous behavior sets in (past the dashed line), where the cell number increases essentially continuously. **(b)** The time derivative of *N* (*t*) (growth rate) exhibits an oscillatory pattern where peaks indicate rapid cell division events and valleys represent time points where divisions do not occur. 5 clear peaks are visible. Peak width (see inset) is a measure of the heterogeneity in the cell cycle times, extracted from the full width at half maximum (FWHM). **(c)** Heterogeneity in the cell cycle time extracted from the FWHM in **(b)** increases with cell cycle number. The spread in cell division time is better fit with the exponent of 1*/*1.6 higher than the predicted 1*/*2 dependence.

We focus on two distinctive features. First, in the initial regime (100 *min* ≤ *t* ≤ 185 *min*), *N* (*t*) exhibits clear step-like patterns (see the red arrows in Fig. 1a). The time intervals, with constant *N*, are interspersed with time regimes with increasing *N* (see the shaded regions). The step like patterns are a direct consequence of synchronous cell divisions, punctuated by finite time intervals where there are no cell division events. Second, at later times (*t* ≥ 185 *min*), the staircase pattern in *N* (*t*) disappears, signaling the onset of the synchronous to asynchronous transition (SAT), after 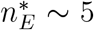^*th*^ round of cell division. Here, the subscript “E” denotes experiments. These peculiar patterns raise the following questions:what sets the time scale for SAT? Does that provide clues as to the mechanism behind the SAT? which motivated us to develop the theoretical framework discussed below.

## Increasing fluctuations in cell division times drive the SAT

It is evident from the experimental data^18^ in Fig. 1a-b that the SAT occurs after 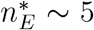 rounds of cell division. Our objective is to provide a theory to predict the number of cell division rounds, 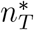 (*T* stands for theory), it takes for SAT. Let the *n*^*th*^ round of division for an individual cell occur at time, *t*_*n*_, which we consider to be a random variable,

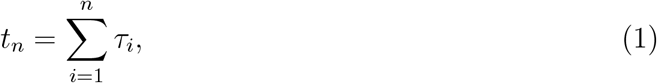

where *τ*_*i*_ is the cell cycle time for the *i*^*th*^ round of cell division (see Fig. 2 in the SI). We assume that all the *τ*_*i*_’s are independent and identically distributed random numbers with the statistics of *τ*_*i*_’s being given by 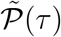 as derived in section I in the SI. From Eqn.(1), we can readily obtain the statistics of *t*_*n*_. We will focus on the mean 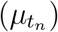 and the variance 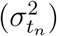 because they can be compared to the experimental data^18,20^. The mean is given by

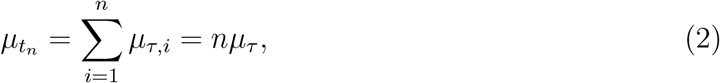

and the variance

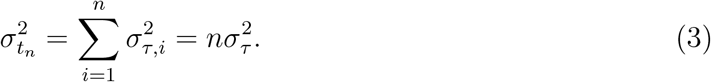

**FIG. 2:**
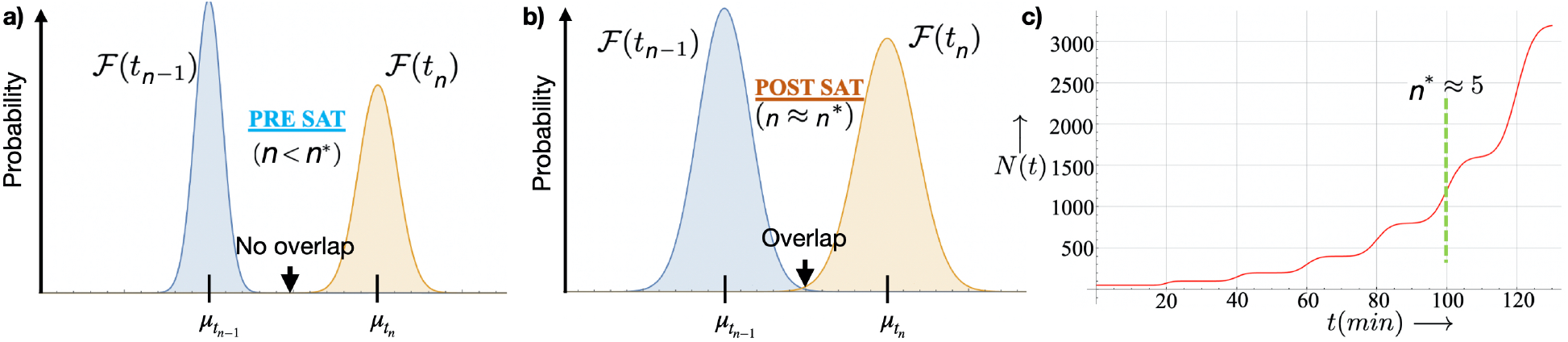
Variation in cell division times. **(a)** Theory to predict the onset of the SAT. Pre-SAT: the probability distributions, ℱ(*t*_*n*−1_) and ℱ (*t*_*n*_) of the times at which cells divide, do not overlap, implying that cell divisions do not occur for a finite time regime between *t*_*n*−1_ and *t*_*n*−1_. **(b)** Post-SAT: we propose that ℱ (*t*_*n*−1_) and ℱ(*t*_*n*_) begin to overlap, with cell divisions throughout a given time interval. **(c)** The plot of the number of cells (*N* (*t*)) for multiple cell division rounds (*n*) shows a staircase-like pattern, using theory. The dashed line indicates the onset of SAT at *n*^*^ ≈ 5. We reproduce the experimentally observed *N* (*t*) curve during Zebrafish morphogenesis and the number of cell division cycles necessary for the onset of SAT^19^.

Here, *µ*_*τ*_ and *σ*_*τ*_ are the mean and standard deviation of a single cell cycle round (see SI section I for details). Both the mean and the variance increase linearly with the number of cell cycle rounds (*n*) as seen from Eqs. (2) and (3). We predict that the probability distribution for *t*_*n*_ is given by

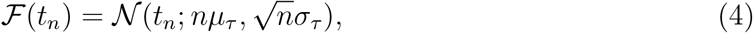

where 𝒩 is a normal distribution with mean *nµ*_*τ*_, and standard deviation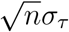.

To test the theoretical predictions, we quantified the statistics of cell cycle times from the experimental data. If our model is correct, we expect the fluctuations in cell division times to increase at later cell division cycles. The shaded regions in Fig. 1a mark the time intervals where divisions occur and visibly broaden with time. However, a concrete way to quantify the fluctuations in cell division time is needed. Calculating the number growth rate 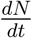 allows us to extract the spread in cell division times as a function of *n* from the full width at half maximum (FWHM) of each peak (see Fig. 1b and inset). Remarkably, as predicted by theory (see Fig. 1) the variance in cell division times increase with *n*. The spread in cell division times scales as 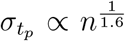, close to the theoretical prediction 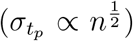.The modest deviation from theory might be due to limited statistics, warranting further targeted experiments to test the functional dependence of fluctuations with time. We note that Eq. (4) can be interpreted as a particle undergoing Brownian motion advected by a constant “velocity” *µ*_*τ*_. Although variability in the cell cycle duration was observed long ago, its origin is unclear^21,22^. Our theory suggests that the fluctuations in cell division times has an important consequence in driving SAT.

With the ansatz for ℱ (*t*_*n*_), we can predict the number of cell division cycles (*n*^*^) where SAT occurs. Generally, we define the onset of SAT to be the absence of a plateau in the *N* (*t*) curves between successive cell division epochs. This is realized when the fluctuations in *t*_*n*\*_ are so large that the cell division time probabilities at two consecutive cell cycles – 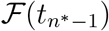 and 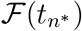 - begin to overlap (see Figure 2a and 2b). Mathematically, this is equivalent to writing,

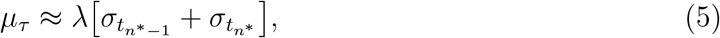

where *λ* controls the strength of the overlap between two successive distributions, ℱ’s. We assume that the two distributions start to overlap at three standard deviations from the mean, *λ* = 3 (see Fig. 2b). Note that *λ* can vary depending on the strength of the overlap between the ℱ’s. Using *λ* = 3, we calculated *n*^*^, the cell cycle round when SAT occurs as,

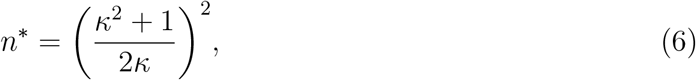

where 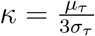. In the experiments, *σ*_*τ*_ = 1.6 *min* and *µ*_*τ*_ = 20 *min*. Therefore, *n*^*^ = 4.85 ≈ 5 from Eqn. (6), close to the experimental data in Figure 1a, where the transition occurs after five cell division rounds. See SI Figure 3 for an analysis on how *n*^*^ depends on *λ*. The theory also predicts that at the transition, 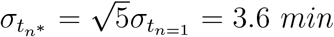, which is close to the experimentally observed value of 2.8 *min*. The choice of *λ* = 3 is justified *a posteriori*.

**FIG. 3:**
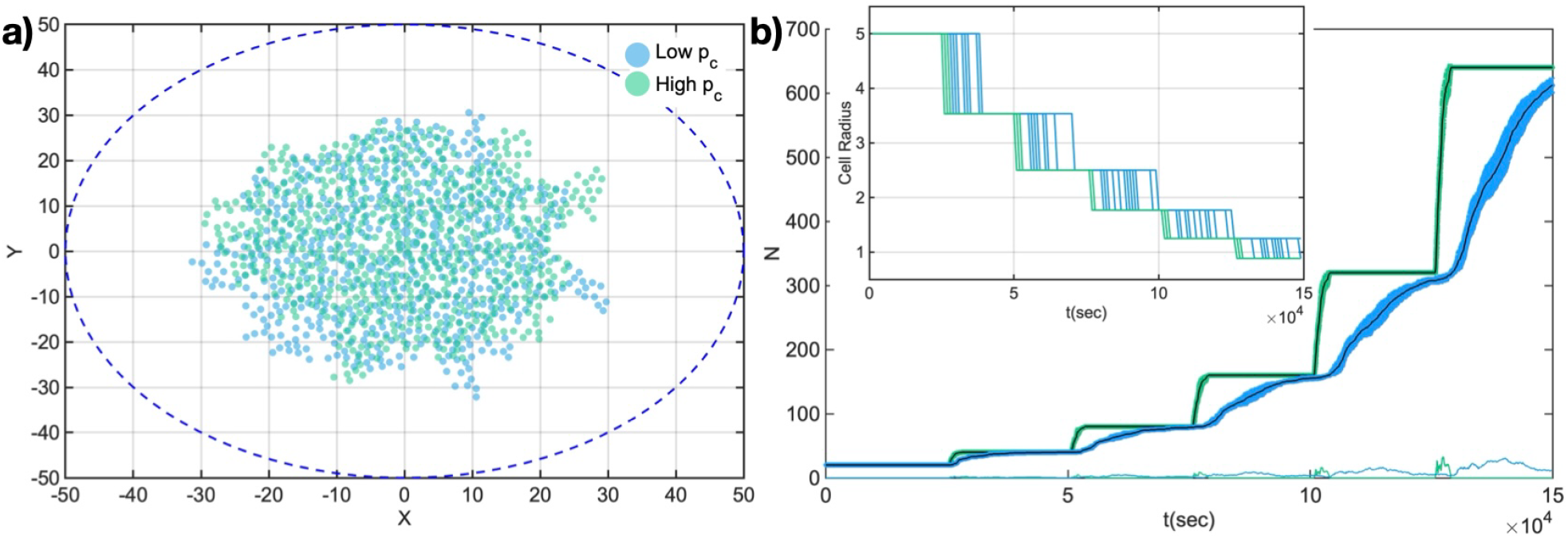
Mechanical feedback determines synchronous cell division. **(a)** Snapshots of the cell collective at *t* = 150, 000 *s* for two different simulation runs at low *p*_*c*_ = 1.0 × 10^−5^*MPa* and high *p*_*c*_ = 1.0 × 10^−4^*MPa*. The blue dashed line denotes the boundary. **(b)** Number of cells vs time shows that at low *p*_*c*_ cell divisions are rapidly desynchronized as opposed to high *p*_*c*_. Mean values of *N* (*t*) is shown in black and shaded region is the standard deviation (over 3 simulation runs). We show the standard deviation as a function of time at the bottom. Inset: plot of cell radii vs time showing that cells decrease in size with each successive division cycle closely mimicking *in vivo* observations.

## Step-like pattern in cell number growth is recapitulated in theory

Our theory captures the step-like pattern in the growth of *N* (*t*) observed in experiments (see Figure 1a). We estimate that *N* (*n, t*) which denotes the number of cells at time *t* after *n* rounds of cell division from the following relation,

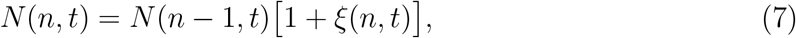

where *ξ*(*n, t*) is the probability that a cell has undergone *n* rounds of cell division until time *t*. In particular, *ξ*(*n, t*) is the cumulative distribution function (CDF) associated with ℱ (*t*_*n*_) and is given by,

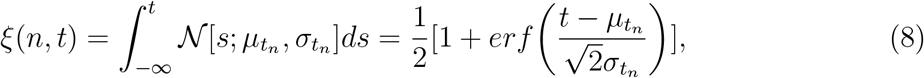

where *erf* is the error function. In the above equation, the lower integration limit is from −∞ because the support of the normal distribution is from [−∞, ∞]. However, since the standard deviation, 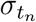, is small compared to 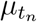, the majority of the contribution to integral in Eq. 8 arises from 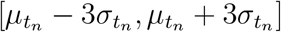.Substituting the expression for *ξ*(*n, t*) into Eq. 7, we obtain the recursive relation,

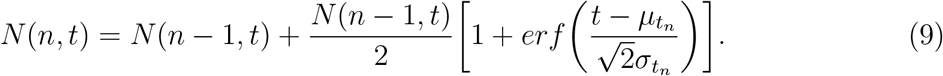

Given the initial condition, *N* (*n* = 0, *t* = 0), we can calculate *N* (*n, t*) for any *n* and *t*. Remarkably, the plot for *N* (*n, t*) using Eqn. (9) reproduces the step-like pattern in cell number growth (compare Figure 1a and Figure 2c).

Although our theory shows that synchronous cell division can be quantitatively described without considering interactions between cells, the onset of asynchronous cell division together with increasing cell numbers, may point to the importance of the local physical forces experienced by cells in controlling division^17,23,24^. Seminal studies have shown that variations in cell density can control cell proliferation rate through contact inhibition of proliferation^25^. Contact inhibition restricts cell proliferation when space is limited as dense cell spatial arrangements lead to strong suppression of cell divisions^26,27^. To determine whether and how intercellular mechanical interactions influence SAT, we implement a two-dimensional (2D) agent-based model for cell collectives characterized by mechanical feedback-dependent cell divisions.

## Biomechanical feedback controls the SAT

Regulating the cell cycle based on mechanical feedback could have significant implications, especially in cancer therapy. As tumors generate physical forces during growth^28^, a feedback circuit to control tumor cell divisions due to growth generated forces could be the basis for alternative therapeutic approaches.

To elucidate how biomechanical feedback impacts cell division, we start with the hypothesis that cell-cell contact leads to forces that limit a cell’s ability to progress through the cell cycle. The cumulative effect of the forces experienced by a cell from its microenvironment is encoded through the pressure on cells which we predict will decrease the rate of division^17,20,24^. To test the consequences of such mechanical feedback on cell division, we performed agent-based simulations built on our earlier works.^24,29–34^ The basic rule that we consider for cell cycle progression is a “timer mechanism” such that a cell divides after the evolution of a fixed time, *τ*_*min*_. Cells can only progress through the cycle and divide if the pressure they experience is below a critical pressure (*p < p*_*c*_).

To account for experimental observations^7^, we assume that cells that do not grow in size^35^. Conversely, after each cell division, daughter cell sizes (*R*_*d*_) are reduced to conserve the total area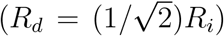. This implies that the cell size decreases gradually with each cell division. Recently, the collective cell dynamics due to cell size reductional division was investigated^36^. We track the sum of the normal pressure, *p*_*i*_, that the *i*^*th*^ cell experiences due to contact with its neighbors (see SI Section III for details on the simulation model). Importantly, lower *p*_*c*_ implies that cells enter the dormant phase with high probability, while higher *p*_*c*_ values ensure that cells are less sensitive to feedback from intercellular mechanical interactions. We predict that higher sensitivity to mechanical feedback will lead to a faster onset of asynchronous cell divisions as the individual cell cycle timer would be easily paused due to mechanical feedback, also causing a broadening of the cell division times.

By implementing these rules for mechanical feedback into a 2D agent-based model, the *N* (*t*) pattern closely matches the experimental observations. The snapshot of the cell collectives under strong (low *p*_*c*_) and weak (high *p*_*c*_) mechanical feedback in Fig. 3a show no marked differences in the overall number and cell spatial distribution. However, clear steps of cell division events followed by time regimes with absence of division are visible at high *p*_*c*_ (see Fig. 3b). As predicted, the synchronous division events persist at higher *p*_*c*_ and are maintained for longer times. On the other hand, at lower *p*_*c*_, cell divisions occur continuously (compare blue and magenta curves in Fig. 3b). The inset shows decreasing cell radii for multiple individual cells at lower and higher values of *p*_*c*_.

Our results imply that cell sensitivity to mechanical feedback is the key determinant of the speed with which the onset of SAT occurs. Cells with low sensitivity to mechanical feedback would undergo synchronous cell division across multiple cell cycles. However, cells that are highly sensitive to mechanical feedback would quickly transition to asynchronous cell division cycles. We conducted additional 3D simulations which confirm that mechanical feedback controls the onset of SAT (see SI section IV). Even with cell size growth that we consider in the 3D model, our conclusions remain unchanged.

## Mechanical feedback lengthens the cell cycle

So far we have focused on the collective effect of coordinated cell division events between cells. We now focus on the single cell scale which further lends support to our prediction that mechanical feedback lengthens the cell cycle. Agent-based modeling is a powerful approach to bridge scales and allow us to relate emergent properties at the cell collective scale to the individual cellular behaviors. Specifically, the rules we implemented for timer based cell cycle progression will have consequences on how mechanical pressure affects the cell cycle times. Many embryos exhibit a period of rapid cell division followed by lengthening cell division times during early development^37^. The embryo development processes are crucially related to the speed of cell division and the cell cytoskeleton^37,38^ as morphogenetic events require cytoskeletal arrangements, which must wait for cell divisions to slow^37^. Thus, slowing cell division time is an important and conserved feature of embryo development.

We anticipate that pressure-dependent feedback on cell division will lead to the lengthening of the cell cycle. Recently, we showed that there are emergent variations in the cell division rate between the core and periphery of multicellular spheroids due to pressure differences ^24,30,39^. Hence, we turned to our simulations to quantify the effect of pressure on the cell cycle time. Because cells decrease in size with each division, we can quantify the time it takes for individual cells to divide from cell radius vs. time (see the inset of Fig. 3b). The time interval between two subsequent drops in cell radii is the effective time it takes for a cell to divide. The effective division times are longer at low *p*_*c*_ as compared to high *p*_*c*_ (see inset of Fig. 3b). The probability distribution of cell division times (see Fig 4a and inset) confirm this observation, as cell cycle times are longer and much more variable at lower *p*_*c*_.

**FIG. 4:**
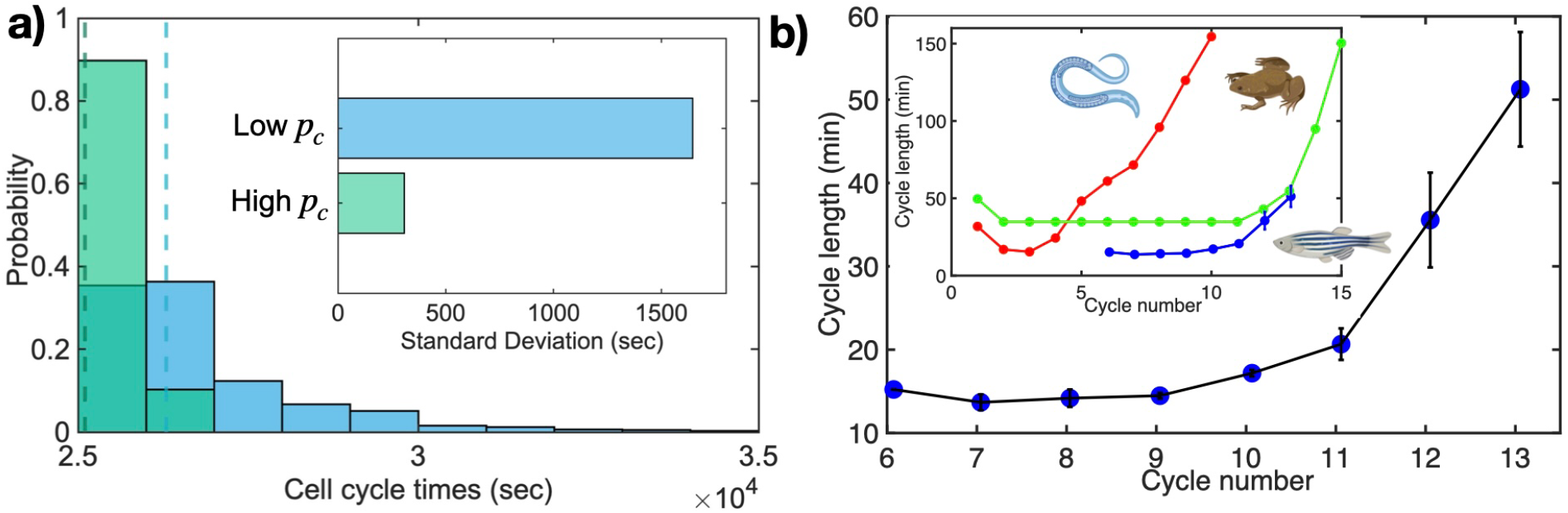
Mechanical feedback elongates and increases the variability of cell cycle length. **(a)** Average cell cycle length increases with stronger mechanical feedback. Average cell division time at low *p*_*c*_ is 2.6292 × 10^4^*s* and 2.5103 × 10^4^*s* at high *p*_*c*_ (see dashed lines). Higher *p*_*c*_, corresponding to lesser cell sensitivity to mechanical feedback on cell division, leads to cell division times closer to the set cell cycle time of *τ*_*min*_ = 2.500 × 10^4^*s*. Inset shows the standard deviation is cell division times across ∼ 900 cells. **(b)** Experimental data extracted from Ref.^7^ for cell cycle length during cycle numbers 6-13 for Zebrafish development. Inset shows similar behavior that is conserved across multiple species - *C. elegans* (roundworm), *X. laevis* (frog) in addition to *D. rerio* (zebrafish).

We then looked at experimental data to assess whether cell cycle lengthening is observed during Zebrafish development. As shown in Fig. 4b, the cell cycle lengthens markedly as cells progress through repeated division cycles. Experimental data, therefore, confirms that there is an increase in the mean cell cycle time^19^ together with SAT during Zebrafish morphogenesis. This data is well fit using 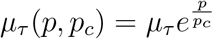, where the exponential increase in cell cycle times is a consequence of mechanical feedback.

## Conclusion

The finding that synchronous cell division depends on the strength of mechanical feedback points to an interesting manifestation of the cross-talk between mechanical forces and cell cycle regulation during embryo development. We show that synchronized cell division is more persistent when mechanical feedback is weaker (i.e. high *p*_*c*_). In contrast, cells under strong mechanical feedback transition rapidly into asynchronous cell division. This implies that suppression of mechanical feedback, which may be realized by partial knockout of E-Cadherin^24^, would prolong the synchronous phase. Another important consequence of mechanical feedback is the lengthening of the cell cycle time. Cells experiencing strong mechanical feedback elongate their cell cycle while cells under weak mechanical feedback progress through the cell cycle more rapidly and more uniformly. The proposed feedback mechanism on growth is therefore a ‘mechanical’ intercellular signal that enables cells to measure density and the timing of development.

## Acknowledgements

We are grateful to Xin Li for useful discussions. AMK acknowledges support from the College of Science and Mathematics at Augusta University. This work was supported by a grant from the National Science Foundation (CHE 2310639) and the Welch Foundation through the Collie-Welch Chair (F-0019).

## Notes

### Competing Interest Statement

The authors have declared no competing interest.

### Summary of Updates

We now consider the scenario of no size growth of cells.

