## Supplementary Information for "Mechanical feedback drives asynchronous cell divisions during embryogenesis"



(Dated: 3 September 2025)



[a] Electronic mail:

[b] Electronic mail:

[c] Electronic mail:

### I. DISTRIBUTION FOR SINGLE CELL CYCLE DIVISION TIMES

The early embryonic cell cycles lack the layers of regulation that is present in more mature cells[1]. To understand the role of mechanical feedback on cell cycle regulation, we introduce a minimal model. The multiple cell cycle stages, shown in Fig.S1a, are represented as discrete steps. Once a cell goes over all the stages, a cell undergoes division. Assuming that there are $n$ stages, let us denote the $i^{th}$ stage of a cell as $G_i$. We denote transition rate from $G_{n-1} \rightarrow G_n$ as $k_n$. We assume, for simplicity, that $k_1 = k_2 = ....k_{n-1} = k$. The functional dependence of $k$ on mechanical feedback ($p$) is given by,

$$k = \left\{ \begin{array}{ll} k & p \leq p_c \\ ke^{-\frac{p}{p_c}} & p > p_c \end{array} \right\}, \tag{1}$$

where $p$ is local mechanical stress on a cell, and $p_c$ is the stress threshold value above which the cell is dormant, at least temporarily (see Fig. S1b). The functional form of $k$ is inspired from experiments, where the cell division time increases as a function of tissue growth[2,3]. For our model, the time ($\tau$) it takes for a single cell to divide is given by,

$$\tau = \sum_{i=1}^{n-1} t_i, \tag{2}$$

where $t_i$ is the transition time from $G_{i-1}$ to $G_i$. Because the $t_i$'s are independent random variables, it follows that $\tau$ is a random variable as well. The assumption that $k_i = k$ for all $i's$ implies that the probability density of $t_i$'s, $\mathcal{P}(t_i)$, are identical, $\mathcal{P}(t_i) = \mathcal{P}(t)$. The probability density, $\mathcal{P}(t)$, is governed by the differential equation,

$$\frac{d\mathcal{P}(t)}{dt} = -k\mathcal{P}(t), \tag{3}$$

that is subject to the constraint $\int_0^\infty \mathcal{P}(t)dt = 1$. The solution to the above equation is,

$$\mathcal{P}(t) = ke^{-kt}. \tag{4}$$

By knowing the functional form of $\mathcal{P}(t)$, we can readily calculate all the moments of $t$. However, in this work, we focus on the mean and the variance as they are the relevant observables in the experiment[4]. The mean, $\mu_t = \langle t \rangle$, and the variance, $\sigma_t^2 = \langle t^2 \rangle - \langle t \rangle^2$, of $\mathcal{P}(t)$ are given as $\frac{1}{k}$ and $\frac{1}{k^2}$ respectively. Utilizing Eq. 2, we can calculate the mean ($\mu_\tau$) of the cell division time, $\tau$, which is given by

$$\mu_\tau = \sum_{i=1}^{n} \mu_t = \frac{n}{k}, \tag{5}$$

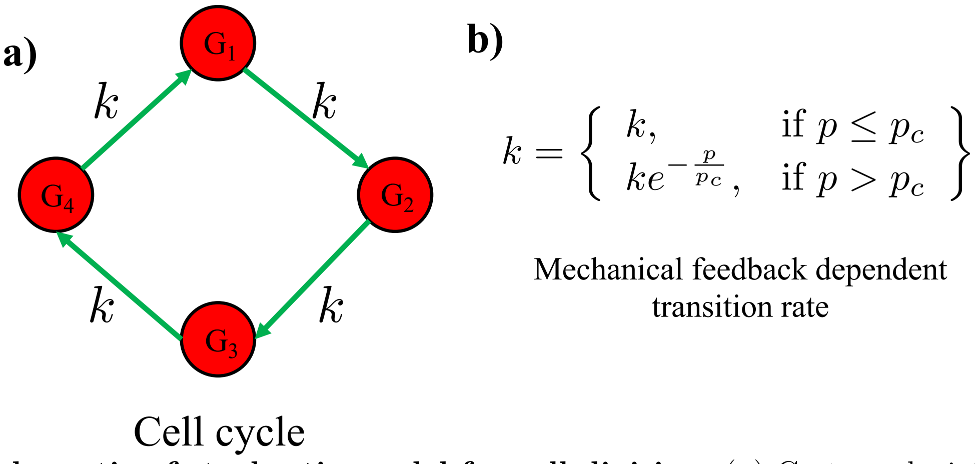


FIG. 1: **Schematic of stochastic model for cell division**. (**a**) Cartoon depicting a cell cycle for an individual cell. In the present picture, the cell cycle is assumed to be comprised of $n = 4$ stages $(G_1, G_2, G_3, G_4)$. A single cell transitions from one stage to the other at the same rate, $k$. Once a cell has gone through all the stages, a cell is assumed to have undergone a single cell division. (**b**) The transition rate between the stages depends on mechanical feedback which is controlled by the local environment (see the expression for $k$ given above.

and the variance $(\sigma_\tau^2)$ of $\tau$ is

$$\sigma_\tau^2 = \sum_{i=1}^{n} \sigma_t^2 = \frac{n}{k^2}. \tag{6}$$

For zebrafish, $\mu_\tau \approx 20\ min$ and $\sigma_\tau \approx 1.6\ min$[4].

In order to analyze the experimental data, we fit the cell division times to a normal distribution (see Fig. 2a in the main text). Thus, for our analyses to be consistent with theory, we ought to recover the normal distribution from theory. This is possible in $n \to \infty$ limit, where we obtain $\tilde{\mathcal{P}}(\tau) = \mathcal{N}(\tau; \mu_\tau, \sigma_\tau)$ utilizing the well known central limit theorem. Here, $\tilde{\mathcal{P}}(\tau)$ denotes the probability distribution of cell division times $\tau$, and $\mathcal{N}(\tau; \mu_\tau, \sigma_\tau)$ is the normal distribution with mean $\mu_\tau$ and standard deviation $\sigma_\tau$. Note that $\tilde{\mathcal{P}}(\tau)$ is the distribution of time, $\tau$, for a cell to undergo a single cell division event.

Physiologically, $n \to \infty$ implies that for a cell to divide, numerous physio-chemical events have to be completed, which occurs with astonishing regularity in many organisms.

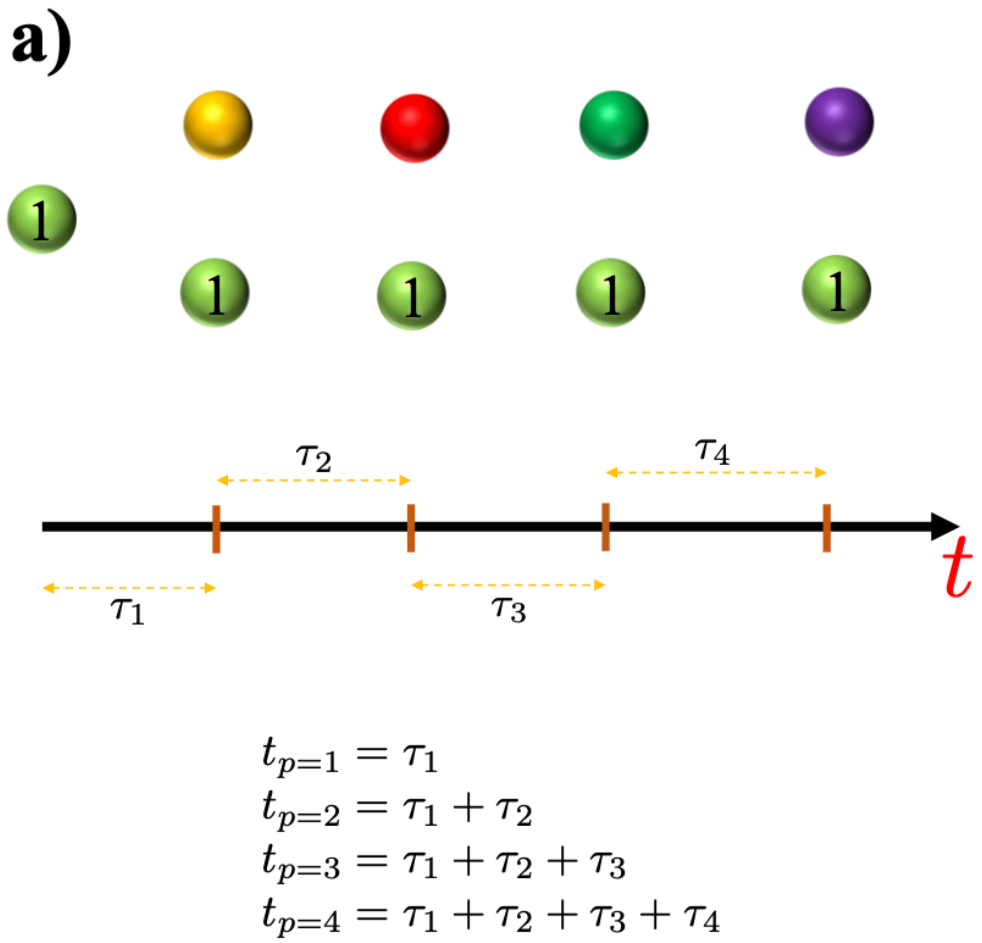


FIG. 2: **Schematic of stochastic model for cell division**. The schematic for a cell (in green labelled as 1) undergoing subsequent cell divisions. The different colored cells at different time points are daughter cells. In the middle, a schematic for Eqn. 1 (main text) is shown. In the bottom, Eq. (1) (main text) is shown mathematically for different $t_p$, where $p$ is the number of cell division rounds.

### II. CHOICE OF $\lambda$

In Eq. 5 of the main text, we need the value of $\lambda$ to predict the transition point. The parameter $\lambda$ sets the degree of overlap between two cell division time probability distributions - $\mathcal{F}(t_{n^*-1})$ and $\mathcal{F}(t_{n^*})$. Figure 3 shows the plot of $n^*$ as a function of $\lambda$. Approximately around $\lambda = 3$, we get the right fit between experiments and theory (see the main text).

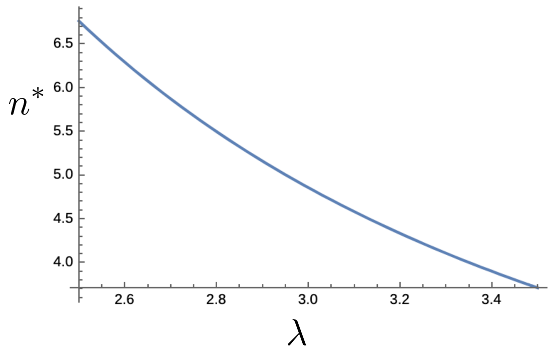


FIG. 3: Plot showing the dependence of $n^*$ on $\lambda$.

### III. 2D SIMULATION DETAIL WITH NO CELL SIZE GROWTH

**Initial Conditions:** At time, $t = 0$, we begin with 20 cells within a circular area of radius of 15 $\mu m$. The radii of the cells are sampled from a Gaussian distribution, $p(R_i) = \frac{1}{0.5\sqrt{2\pi}} e^{-(R_i - 5.000)^2/2\times 0.001^2}$. Similarly, the elastic moduli, $E_i$ and the Poisson ratio $\nu_i$ are obtained from a Gaussian distribution with standard deviation of $10^{-4}$MPa and 0.02 respectively (the mean values are given in Table 1). The receptor and ligand concentration on the cell surface are Gaussian distributed ($p(c_i^{rec}/c_i^{lig}) = \frac{1}{0.02\sqrt{2\pi}} e^{-(c_i^{rec}/c_i^{lig} - 1.00)^2/2\times 0.02^2}$), centered around the mean (=1.0) with a dispersion of 0.02. In subsequent cell division cycles, the daughter cell properties are sampled from the same distribution as described above. At each time step, the growth rate of the cell is also chosen from a Gaussian distribution.

**Boundary conditions:** We incorporated a repulsive force exerted on the cell by a circular wall using a Hookean interaction model. Given the cell's position $\vec{r}_i$, we first compute the radial distance from the origin:

$$O_i = \sqrt{x_i^2 + y_i^2}. \tag{7}$$

If the cell is within the interaction range, meaning its radius $R_i$ plus its position distance $O_i$ exceeds the wall radius $R_w$, a repulsive force is applied. The overlap $\delta_i$ between a cell and the wall is given by:

$$\delta_i = (R_i + O_i) - R_w. \tag{8}$$

The resulting Hookean repulsive force is then computed as:

$$\mathbf{F}_{rep,i} = -k\delta_i\hat{\mathbf{r}}_w \tag{9}$$

where $k$ is the elastic constant and $\hat{\mathbf{r}}_w$ is the unit vector pointing outward from the wall. If no overlap ($\delta_i \leq 0$) is detected, the repulsive force is zero.

**Cell division with no size growth:** We model cell division based on a time-dependent criterion together with a pressure constraint. Division occurs when a cell's internal timer reaches a specific threshold, set to:

$$\tau_{min} = \chi \times 50,000\text{sec} \tag{10}$$

where we can vary the cell division time using the factor $\chi$. In this simulation, we fix $\chi = 0.05$.

Each cell starts with an initial timer value of 1. During each time step, we iterate over all cells. The following conditions govern the cell's fate:

**Growth Phase:** If the cell's timer has not reached the division threshold and the cell is under low pressure ($p_i < p_c$), the timer is incremented indicating progress towards division event.

**Cell Division:** When the timer reaches a multiple of $\tau_{min}$ and the cell is still under low pressure, division occurs. The division process follows these steps:

- The radii of the daughter cells are updated based on a division factor to ensure the total area is preserved upon cell division:

$$R_{\text{new}} = \left(2^{-\frac{1}{2}}\right) R_i \tag{11}$$

- Cell mechanical properties, including modulus, Poisson ratio, receptor density, and ligand density, are assigned to the daughter cell using a random Gaussian distribution.

- The new cell's spatial coordinates are determined by introducing a small displacement in a random angular direction:

$$x_{\text{new}} = x_{\text{parent}} + r_{\text{mitosis}}(1 - 2^{-1/2})\cos(\theta) \tag{12}$$

$$y_{\text{new}} = y_{\text{parent}} + r_{\text{mitosis}}(1 - 2^{-1/2})\sin(\theta) \tag{13}$$

where $\theta$ is a randomly generated angle. The parent cell is also slightly repositioned to ensure proper separation.

Table I lists the parameters used in the simulations.

| **Parameters** | **Values** | **References** |
|---|---|---|
| Timestep ($\Delta t$) | 10s | This paper |
| Friction coefficient ($\frac{\gamma_i}{R_i}$) | 0.0942 kg/($\mu$m s) | This paper |
| Cell Cycle Time ($\tau_{min}$) | 25000 s | [5] |
| Adhesive Coefficient ($f^{ad}$) | $10^{-4}\mu$N/$\mu$m | This paper |
| Mean Cell Elastic Modulus ($E_i$) | $10^{-3}$MPa | [6,7] |
| Mean Cell Poisson Ratio ($\nu_i$) | 0.5 | [7,8] |
| Death Rate ($k_a$) | $10^{-6}$s$^{-1}$ | [7] |
| Mean Receptor Concentration ($c^{rec}$) | 1.0 (Normalized) | [7] |
| Mean Ligand Concentration ($c^{lig}$) | 1.0 (Normalized) | [7] |
| Threshold Pressue ($p_c$) | $0.1 - 1.0 \times 10^{-4}$MPa | This paper |

**Cell-Cell Interactions:** The elastic (repulsive) force between two cells of radii $R_i$ and $R_j$ is modeled as,

$$F_{ij}^{el}(t) = \frac{h_{ij}^{3/2}(t)}{\frac{3}{4}(\frac{1-\nu_i^2}{E_i} + \frac{1-\nu_j^2}{E_j})\sqrt{\frac{1}{R_i(t)} + \frac{1}{R_j(t)}}}, \tag{14}$$

where $E_i$ and $\nu_i$, respectively, are the elastic modulus and Poisson ratio of cell $i$. The overlap between the cells, if they interpenetrate without deformation, is $h_{ij}$, is defined as $\max[0, R_i + R_j - |\vec{r}_i - \vec{r}_j|]$ with $|\vec{r}_i - \vec{r}_j|$ being the center-to-center distance.

Cell adhesion, mediated by receptors on the cell surface, enables the cells to stick together. For simplicity, we assume that the receptor and ligand molecules are evenly distributed on the cell surface. Consequently, the magnitude of the attractive adhesive force, $F_{ij}^{ad}$, between two cells $i$ and $j$ scale as a function of their contact line segment, $L_{ij}$. Keeping the 3D model as a guide[7], we calculate $F_{ij}^{ad}$ using,

$$F_{ij}^{ad} = L_{ij} f^{ad} \frac{1}{2}(c_i^{rec} c_j^{lig} + c_j^{rec} c_i^{lig}), \tag{15}$$

where the $c_i^{rec}$ ($c_i^{lig}$) is the receptor (ligand) concentration (assumed to be normalized to the maximum receptor or ligand concentration so that $0 \leq c_i^{rec}, c_i^{lig} \leq 1$). In the present study, $c_i^{rec}, c_j^{lig}$ are fixed and have been included for consistency with previous studies[7,9,10]. The coupling constant $f^{ad}$ allows us to rescale the adhesion force to account for the variabilities in the maximum densities of the receptor and ligand concentrations. We calculate the contact length, $L_{ij}$, using the length of contact between two intersecting circles, $L_{ij} = \frac{\sqrt{(|4r_{ij}^2 R_i^2 - (r_{ij}^2 - R_j^2 + R_i^2)^2|)}}{r_{ij}}$. Here, $r_{ij}$ is the distance between cells $i$ and $j$. As before, $R_i$ and $R_j$ denote the radius of cell $i$ and $j$.

The the sum of the repulsive and adhesive forces in Eqs.(14) and (23) point along the unit vector $\mathbf{n}_{ij}$ from the center of cell $j$ to the center of cell $i$. The total force on the $i^{th}$ cell is given by the sum over its nearest neighbors ($NN(i)$),

$$\mathbf{F}_i = \Sigma_{j\epsilon NN(i)} (F_{ij}^{el} - F_{ij}^{ad}) \mathbf{n}_{ij}. \tag{16}$$

The nearest neighbors satisfy the condition $R_i + R_j - |\mathbf{r}_i - \mathbf{r}_j| \ > \ 0$.

**Equations of Motion:** If inertial effects are negligible[7], the equation of motion of the $i^{th}$ cell is,

$$\dot{\vec{r}_i} = \frac{\vec{F}_i}{\gamma_i}, \tag{17}$$

where, $\gamma_i = \gamma_i^{\alpha'\beta',visc} + \gamma_i^{\alpha'\beta',ad}$ is the total friction coefficient. The cell-to-matrix friction is given by,

$$\gamma_i^{\alpha'\beta',visc} = 6\pi\eta R_i \delta^{\alpha'\beta'}. \tag{18}$$

The cell-to-cell damping coefficient is,

$$\begin{aligned}\gamma_i^{\alpha'\beta',ad} = \; & \gamma^{max} \Sigma_{j\epsilon NN(i)} (A_{ij} \frac{1}{2} (1 + \frac{\vec{F}_i \cdot \vec{n}_{ij}}{|\vec{F}_i|}) \times \\ & \frac{1}{2} (c_i^{rec} c_j^{lig} + c_j^{rec} c_i^{lig})) \delta^{\alpha'\beta'} .\end{aligned} \tag{19}$$

The indices $\alpha'$ and $\beta'$ represent cartesian co-ordinates. Viscosity of the medium surrounding the cell is denoted by $\eta$ and $\gamma^{max}$ is the adhesive friction coefficient.

**Mechanical feedback through pressure:** We implement a mechanical feedback on cell cycle progression and division on the basis of the local forces experienced by a cell. Depending on the local forces, a cell can either be in the dormant ($D$) or in the growth ($G$) phase. The effect of the local cellular microenvironment on proliferation, a collective cell mechanical effect, is taken into account through the pressure experienced by the cell ($p_i$). We refer to $p_i$ as pressure because it has the same dimensions, and models the mechanical sensitivity of cell proliferation to the local environment. Using Irving-Kirkwood's definition, we calculate the pressure ($p_i$) on the $i^{th}$ cell due to contact with its neighbors[11] using,

$$p_i = \frac{1}{2}\Sigma_{j\epsilon NN(i)}\frac{\mathbf{F}_{ij}\cdot\mathbf{dr}_{ij}}{A_i}, \tag{20}$$

where $A_i$ is local area of influence, equal to $\theta\pi R_i^2$. The proportionality constant $\theta$ serves as a measure to sample the local area around the $i^{th}$ cell and was chosen to be 2. The non-zero value of $p_c$ is the source of biomechanical feedback that regulates cell cycle progression. If the local pressure on the $i^{th}$ cell, $p_i$, exceeds a critical value ($p_c$) the cell immediately ceases to progress through the cell cycle and enters the dormant phase. Note that the cell can progress towards division once $\frac{p_i(t)}{p_c} < 1$. The critical pressure, $p_c$, therefore serves as a mechanical feedback[12]. The local pressure, $p_i$, can easily exceed $p_c$ if it is small. In this case, most cells would be dormant for a long time. In the opposite limit, $p_c \gg p_i$, it is unlikely that the cells will reach the dormant phase. This would result in cell division events. Thus, $p_c$ determines the strength of the mechanical feedback.

There are several ways to compute pressure in cell collectives (for a review see Ref.[13]). Here, we define pressure following previous studies[8,14]. These studies produced quantitative agreement with experiments on the dynamics of multicellular spheroid growth[13].

### IV. 3D SIMULATION DETAILS WITH CELL SIZE GROWTH

**Cell division:** The volume of growing cells increases at a constant rate $r_V$. Cell radii are updated from a Gaussian distribution with the mean rate $\dot{R} = (4\pi R_m^2)^{-1} r_V$, and dispersion of $10^{-7}$. Over the cell cycle time $\tau$,

$$r_V = \frac{2\pi(R_m)^3}{3\tau}, \tag{21}$$

where $R_m$ is the mitotic radius. **Table II lists the parameters used in the 3D simulations.**

**Cell birth, apoptosis and dormancy :** We implement a mechanical feedback on cell growth and division on the basis of the local forces experienced by a cell. Depending on the local forces, a cell can either be in the dormant ($D$) or in the growth ($G$) phase. The effect of the local cellular microenvironment on proliferation, a collective cell mechanical effect, is taken into account through the pressure experienced by the cell ($p_i$). We refer to $p_i$ as pressure because it has the same dimensions, and models the mechanical sensitivity of cell proliferation to the local environment. The total pressure ($p_i$) on the $i^{th}$ cell is,

$$p_i = \sum_{j \in NN(i)} \frac{|F_{ij}|}{A_{ij}}, \tag{22}$$

The non-zero value of $p_c$ is the source of biomechanical feedback that regulates cell growth.

There are several ways to compute pressure in cell collectives (for a review see Ref.[13]). Here, we define pressure following previous studies[8,14]. These studies produced quantitative agreement with experiments on the dynamics of multicellular spheroid growth[13].

**Cell-Cell Interactions:** By building on our previous work[7] and similar to the 2D simulations, we consider the cell-cell interactions to be driven by elastic (repulsive) and adhesive (attractive) forces. The total force on the $i^{th}$ cell is, $\vec{F}_i = \Sigma_{j \epsilon NN(i)}(F_{ij}^{el} - F_{ij}^{ad})\vec{n}_{ij}$, where $F_{ij}^{el}$ and $F_{ij}^{ad}$) are cell-cell elastic and adhesive forces respectively, and $\vec{n}_{ij}$ is the unit vector from the center of cell $j$ to cell $i$. We used the Hertz contact mechanics[8,15,16] to model the elastic forces between the spherical cells. $R_i$ and $R_j$, are the cell radii and the parameters $E_i$ and $\nu_i$, respectively, are the elastic modulus and Poisson ratio of the $i^{th}$ cell in the elastic force term: $F_{ij}^{el} = \frac{h_{ij}^{3/2}}{\frac{3}{4}(\frac{1-\nu_i^2}{E_i} + \frac{1-\nu_j^2}{E_j})\sqrt{\frac{1}{R_i(t)} + \frac{1}{R_j(t)}}}$. The overlap between two cells, $h_{ij}$, is defined as $\max[0, R_i + R_j - |\vec{r}_i - \vec{r}_j|]$.

The inter-cell adhesive force, $F_{ij}^{ad}$, is given by,

$$F_{ij}^{ad} = A_{ij} f^{ad} \frac{1}{2}(c_i^{rec} c_j^{lig} + c_j^{rec} c_i^{lig}). \tag{23}$$

In the above equation, $A_{ij} = \pi h_{ij} R_i R_j / (R_i + R_j)$ is the cell-cell contact area, $c_i^{rec}$ ($c_i^{lig}$) is the adhesion receptor (ligand) concentration (assumed to be normalized with respect to the maximum receptor or ligand concentration such that $0 \leq c_i^{rec}, c_i^{lig} \leq 1$)[8]. The dimensions of the adhesive strength, $f^{ad}$, in Eq. 23 is $\mu N/\mu m^2$.

**Boundary conditions:** We initiated the simulations by generating 50 cells, randomly distributed in a cubic region within a 3D spatial domain. For all future time steps, we consider an open boundary condition.

By implementing these rules for mechanical feedback into a 3D agent-based model, we observe that the growth rate of the cell collective closely recapitulate experimental data where clear peaks with bursts of cell division events are followed by time regimes with absence of division events (see Figs. 4a-c). Notably, the synchronous division events are more persistent at higher $p_c$ and are maintained for longer times as compared to lower $p_c$ where the clear peaks in division rate are quickly washed out (compare Figs. 4a and c). To quantify the synchronous nature of the cell division events, we evaluated the autocorrelation of the division rate to verify the presence of cycles and determine their duration. The autocorrelation function of a periodic signal has the same periodic characteristics, evaluated using,

$$C(t_\gamma) = \langle \Sigma_{\alpha,\beta} \frac{dN(t_\alpha)}{dt} \times \frac{dN(t_\beta)}{dt} \delta(t_\gamma - (t_\alpha - t_\beta)) \rangle, \tag{24}$$

where, $t_\gamma$ is the lag time between time points $t_\alpha, t_\beta$, and $\delta(z) = 1$ if $z = 0$ and 0 otherwise. The autocorrelation function is normalized to ensure that $C(t_\gamma = 0) = 1$. We focus on the first peak in the autocorrelation function in order to quantify the peak height and peak location as a function of $p_c$. The peak height measures the strength of the oscillatory division patterns. The peak height increases and saturates at high $p_c$, as shown in Fig. 4e, which is indicative of more persistent synchronous division patterns with weaker mechanical feedback (see Fig. 4f).

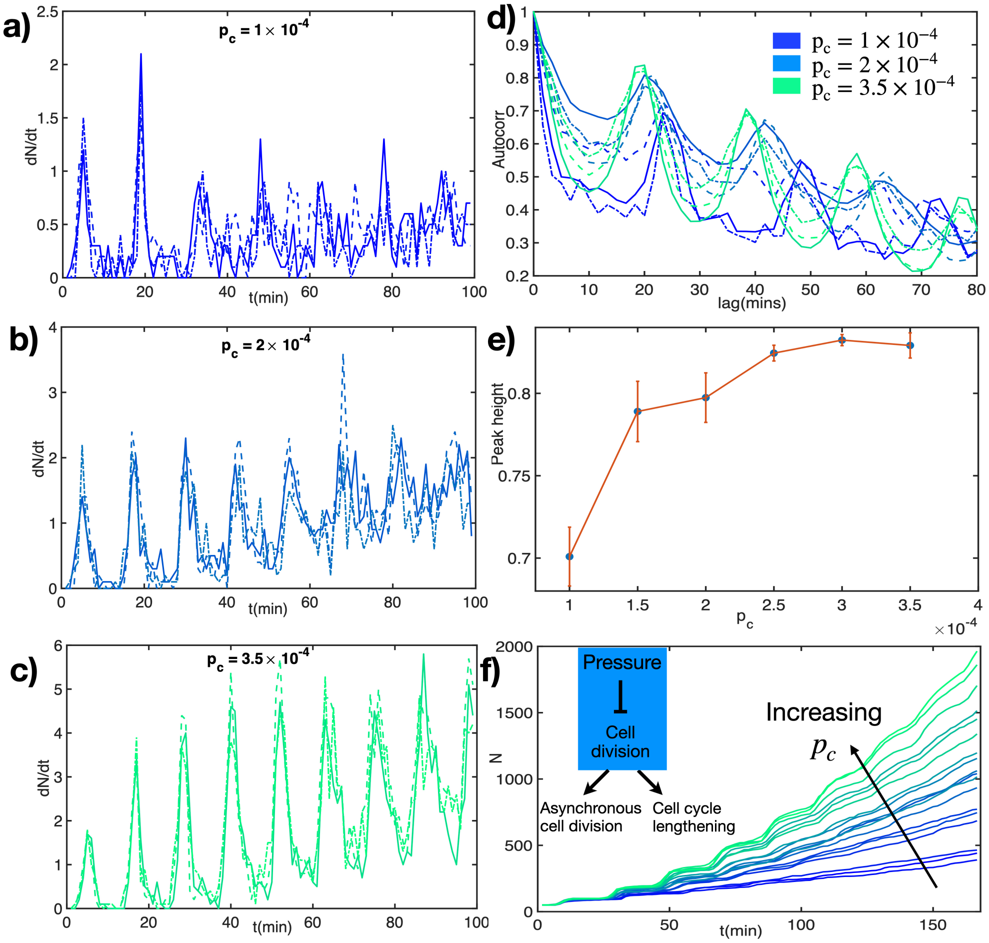


FIG. 4: **Mechanical feedback determines synchronous cell division (a)** Division rate as a function of time at low $p_c = 1 \times 10^{-4} MPa$. Low $p_c$ value corresponds to heightened cell sensitivity to mechanical feedback on cell growth and division. **(b-c)** Division rate as a function of time at intermediate and high $p_c = 2.5 \times 10^{-4} MPa$ and $p_c = 3.5 \times 10^{-4} MPa$, respectively. Higher $p_c$ value, corresponding to lesser cell sensitivity to mechanical feedback on cell growth and division, leads to highly synchronous cell division for a longer time range. **(d)** Autocorrelation of cell division rate at 3 $p_c$ values. The height and time point of the first peak determine the strength of periodicity and the interval between repeated peaks in cell divisions. **(e)** Peak height of the autocorrelation function at varying sensitivity to mechanical feedback through pressure. Strength of periodic division rate increases with $p_c$ and saturates at higher values. **(f)** Number of cells vs time as generated from simulations incorporating mechanical feedback on cell divisions. Inset: Schematic illustrating how increasing pressure limits cell divisions, leading to cell cycle lengthening and asynchronous cell divisions.

Table II: Parameters used in the 3D simulations.

| **Parameters** | **Values** | **References** |
|---|---|---|
| Critical Radius for Division ($R_m$) | 5 $\mu$m | [8] |
| Extracellular Matrix (ECM) Viscosity ($\eta$) | 0.005 kg/($\mu$m s) | [6] |
| Benchmark Cell Cycle Time ($\tau_{min} = k_b^{-1}$) | 900 s | [5] |
| Adhesive Strength ($f^{ad}$) | $1 \times 10^{-4}\mu$N/$\mu$m$^2$ | [8], This paper |
| Mean Cell Elastic Modulus ($E_i$) | $10^{-3}$MPa | [6] |
| Mean Cell Poisson Ratio ($\nu_i$) | 0.5 | [8] |
| Death Rate ($k_a$) | $10^{-6}$s$^{-1}$ | This paper |
| Mean Receptor Concentration ($c^{rec}$) | 0.90 (Normalized) | [8] |
| Mean Ligand Concentration ($c^{lig}$) | 0.90 (Normalized) | [8] |
| Adhesive Friction $\gamma^{max}$ | $10^{-4}$kg/($\mu$m$^2$ s) | This paper |
| Threshold Pressue ($p_c$) | $1.0 - 3.5 \times 10^{-4}$MPa | [8,17] |
